# Statistical learning occurs during practice while high-order rule learning during rest period

**DOI:** 10.1101/2020.10.25.353375

**Authors:** Romain Quentin, Lison Fanuel, Mariann Kiss, Marine Vernet, Teodóra Vékony, Karolina Janacsek, Leonardo Cohen, Dezso Nemeth

**Author notes:** These authors contributed equally to this work.

## Abstract

Knowing when the brain learns is crucial for both the comprehension of memory formation and consolidation, and for developing new training and neurorehabilitation strategies in healthy and patient populations. Recently, a rapid form of offline learning developing during short rest periods has been shown to account for most of procedural learning, leading to the hypothesis that the brain mainly learns during rest between practice periods. Nonetheless, procedural learning has several subcomponents not disentangled in previous studies investigating learning dynamics, such as acquiring the statistical regularities of the task, or else the high-order rules that regulate its organization. Here, we analyzed 506 behavioral sessions of implicit visuomotor deterministic and probabilistic sequence learning tasks, allowing the distinction between general skill learning, statistical learning and high-order rule learning. Our results show that the temporal dynamics of apparently simultaneous learning processes differ. While general skill and high-order rule learning are acquired offline, statistical learning is evidenced online. These findings open new avenues on the short-scale temporal dynamics of learning and memory consolidation and reveal a fundamental distinction between statistical and high-order rule learning, the former benefiting from online evidence accumulation and the latter requiring short rest periods for rapid consolidation.

## Introduction

Learning is the ability to acquire knowledge or skills through new or repeated experiences. To understand the neural mechanisms of learning, it is crucial to identify the specific periods during which it occurs. In the laboratory, learning is usually assessed by measuring specific knowledge or skill before and after a period of training. For example, a seminal experience consists of measuring the speed and accuracy with which participants play a sequence – a simplified version of learning a piece of piano without the artistic component – before and after practicing it several times (Fischer et al. 2002). This type of research revealed that following a training session and during a resting or sleep period, the acquisition of new skill may continue to develop, a process called offline learning (Robertson et al. 2004). Indeed, performance (Robertson et al. 2005) or the stability of the memories against interference (e.g., caused by the learning of a second sequence) (Brashers-Krug et al. 1996; Shadmehr and Brashers-Krug 1997) is enhanced several hours after the end of the practice compared to just after the practice. This offline learning, which occurs during awake or sleep periods, has been linked to functional brain changes (Shadmehr and Holcomb 1997; Fischer et al. 2005). This demonstrates that the neural mechanisms of learning do not necessarily only develop during practice. Recently, rapid offline consolidation of skill has also been documented in the course of short rest periods, from seconds (Bönstrup et al. 2019, 2020) to minutes (Du et al. 2016) during the learning of a perceptual-motor sequence. In Bönstrup et al. (2019, 2020), this fast offline learning even accounted for most behavioral gains during early skill learning, raising the hypothesis that the brain mainly learns during short rest periods and not during the practice itself. However, these studies investigating ultra-fast consolidation during sequence learning did not evaluate the relative contribution of online and offline learning to different crucial components of learning. Here, we used sequence learning tasks with random, probabilistic and deterministic transitions that made possible the identification of the short-scale dynamics of general skill (the general speed-up in the task), statistical and high-order rule learning.

Statistical learning is a fundamental learning mechanism responsible for picking up probabilistic regularities in the environment. The ability of an organism to extract such statistical environmental information is critical for its survival (Saffŕan et al. 1996; Milne et al. 2018) and is present across species and modalities (Bulf et al. 2011). In humans, this ability is present in babies (Saffran et al. 1996) and at the core of a wide range of behaviors, including linguistic processing (Saffran et al. 2001) or perceptual decision making (Summerfield and de Lange 2014). One challenge of language acquisition, for example, is the segmentation of words from fluent speech. Within a language, the transitional probability between two syllables will generally be higher within a word than between two words, creating inhomogeneities in transitional probabilities between sounds. Such statistical information is used by adults and babies as young as 8 months old in order to segment words (Saffran et al. 1996; Mirman et al. 2008).

Nevertheless, learning does not rely solely on the extraction of statistical regularities. High-order rule learning is also needed to extract deterministic rules that can be generalized to new elements that have never been encountered before. For instance, it has been shown that 7-months-old babies can also extract and generalize abstract rules from an artificial language (Marcus et al. 1999) and that these rules are captured during speech processing (Peña et al. 2002). Such rules are abstract in the sense that they can be applied to new elements in the environment that have never been encountered before. They are often said to be “high-order” because the knowledge of several elements (n-1, n-2, etc.) is necessary to predict an upcoming element (n). Well-beyond language acquisition, the brain is constantly making predictions based on previous knowledge in virtually all types of learning (Engel et al. 2001; Friston 2005; Kveraga et al. 2007). Such predictions may be inferred from both statistical regularities and high-order rules. Here, we explore whether statistical learning and high-order rule learning are related to different ultra-fast consolidation dynamics.

Learning a new visuomotor skill also requires the development of lower-level perceptual and motor skills that do not depend on statistical or high-order rule learning, including visuomotor mapping and dexterity (Robertson 2007). We refer to this type of learning as general skill learning.

In this study, we used serial reaction time (SRT, Nissen and Bullemer 1987) and alternating serial reaction time tasks (ASRT, Howard and Howard 1997), in which healthy participants encounter an array of four positions on a screen, each paired with a designated response key. Positions are filled sequentially with deterministic (in both SRT and ASRT) or probabilistic (in ASRT) patterns and the participant has to push the corresponding key as fast and as accurately as possible. These task designs allow the distinction between general skill learning, statistical learning and high-order rule learning. We identified the short-scale temporal dynamics of these three types of learning by measuring the performance gains during short practice (online) or rest (offline) periods. Beyond confirming that general skills are mainly learned during short rest periods (Du et al. 2016; Bönstrup et al. 2019, 2020), our analyses revealed a critical distinction between statistical learning that is acquired during practice and high-order rule learning that is acquired during rest periods. These results suggest that the brain mechanisms leading to statistical and high-order rule learning are fundamentally different, the former requires online evidence accumulation while the latter requires a rest consolidation period.

## Results

### Dynamics of general skill learning

In the three experiments, average RT per block for all trials (excluding random blocks in the SRT task) decreased over time (Experiment 1: Spearman *r_s_* = −0.97, *p* < 10^−12^; Experiment 2: *r_s_*= −0.78, p < 10^−9^; Experiment 3: *r_s_*= −0.99, *p* < 10^−155^), demonstrating general skill learning (Fig. 2A, 2E, 2H for respectively Experiment 1, 2 and 3, black line). To investigate whether this learning occurred during practice or rest periods, we measured its online and offline contribution as depicted in Fig. 1B and described in the methods section. In average across blocks, general skill performance decreased during practice (Experiment 1: *M_online_* = −24.58 ± 22.79 seconds, *t*(62) = −8.49, *p* < 10^−11^; Experiment 2: *M_online_* = −24.23 ± 9.37 seconds, *t*(179) = −34.61, *p* < 10^−80^; Experiment 3: *M_online_* = −13.97 ± 6.63 seconds, *t*(24) = −10.32, *p* < 10^−9^) and increased during rest periods (experiment 1: *M_offline_* = 37.22 ± 24.78 seconds, *t*(62) = 11.88, *p* < 10^−16^; Experiment 2: *M_offline_* = 25.66 ± 9.35 seconds, *t*(179) = 36.71, *p* < 10^−84^; Experiment 3: *M_offline_* = 20.29 ± 26.35 seconds, *t*(24) = 11.59, *p* < 10^−10^) (Fig. 2C, 2F, 2I).

**Figure 1:**
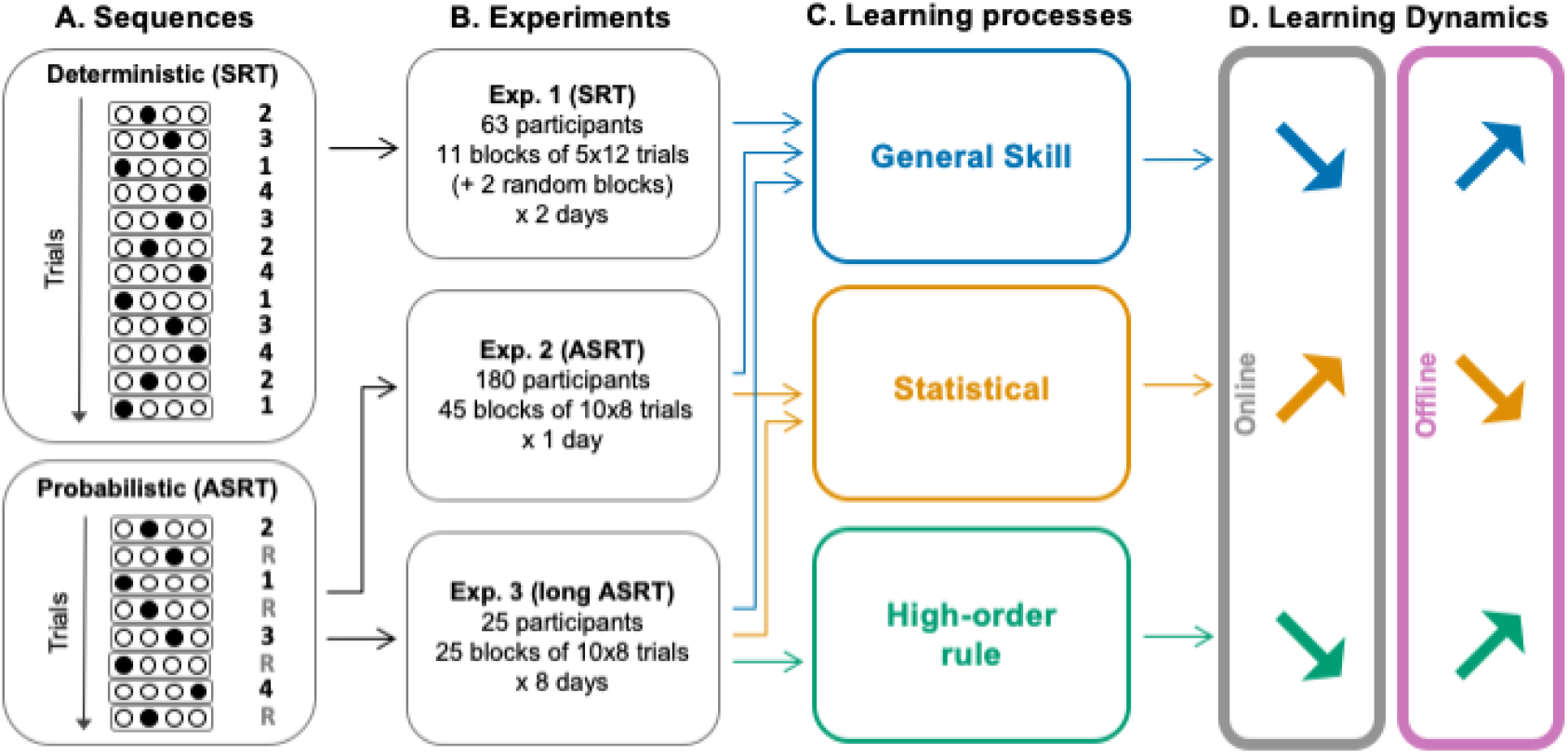
General design and main results. (A) Structure of the sequences used in the SRT task and ASRT task. In the SRT task, a deterministic sequence of 12 elements is repeated five times per block. In the ASRT task, a deterministic sequence of 4 elements is interleaved with four random elements resulting in an 8-element probabilistic sequence, which is repeated ten times per block. (B) The number of participants and sessions in Experiment 1 (SRT experiment), Experiment 2 (ASRT experiment), and Experiment 3 (long ASRT experiment), (C) Type of learning investigated in each experiment. (D) Summary of the results. General skill and high-order rule learning occur during rest periods (offline) while statistical learning occurs during practice (online).

**Figure 2:**
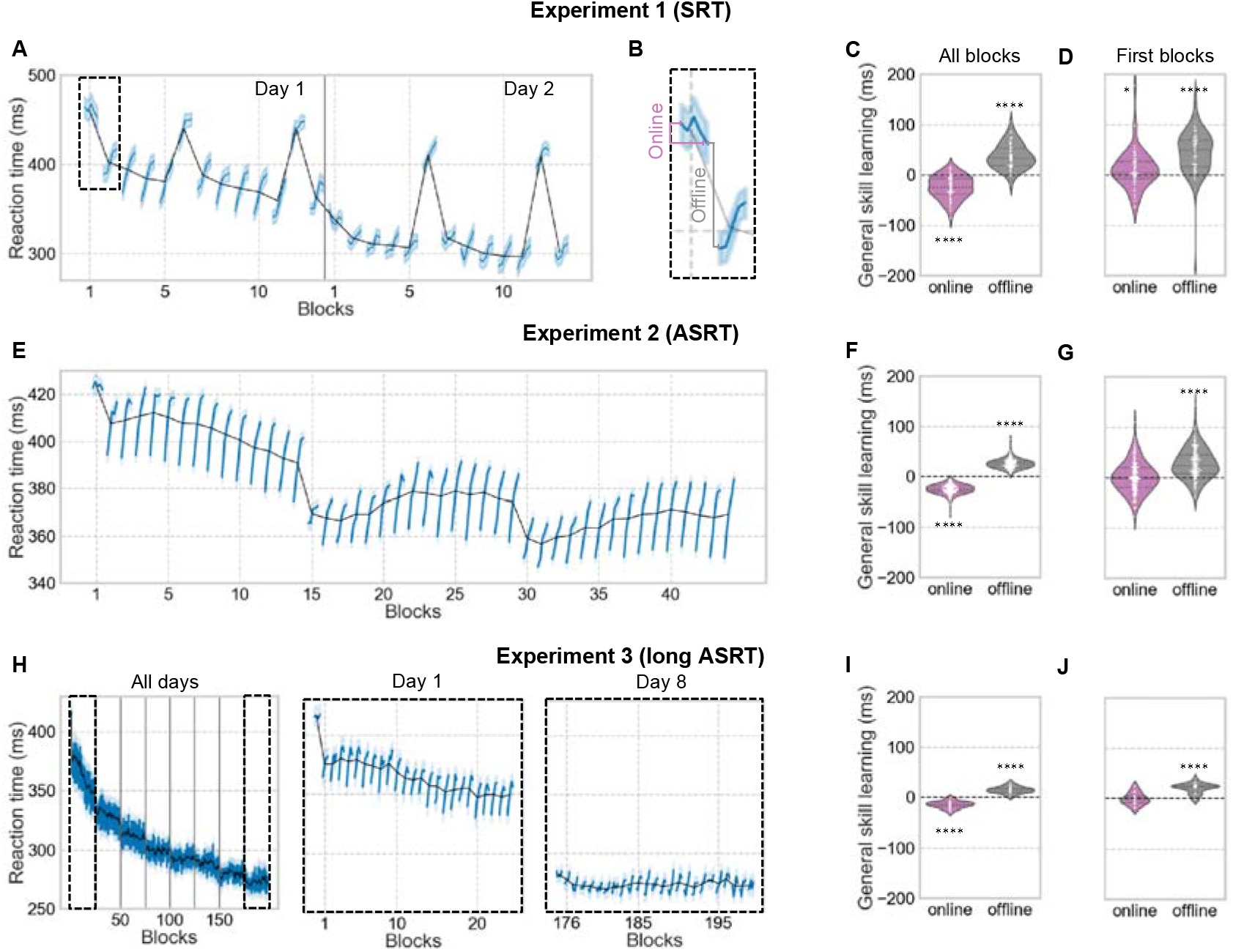
General skill learning occurs during rest periods. (A) Average reaction time per block (black line) and per bin (blue line) for the Experiment 1 (SRT). (B) Depiction of online and offline learning measurement. (C) Average online and offline general skill learning across all blocks and (D)for only the first block of both sessions for Experiment 1 (SRT). (E) Average reaction time per block (black line) and per bin (blue line) for Experiment 2 (ASRT). (F) Average online and offline general skill learning across all blocks and (G) for only the first block for Experiment 2 (ASRT). (H) Average reaction time per block (black line) and per bin (blue line) for Experiment 3 (long ASRT). For better visualization, a zoom-in for day 1 and day 8 is represented. (I) Average online and offline general skill learning across all blocks and (J) for only the first block of the 8 sessions for Experiment 3 (long ASRT). Significance is noted by * for p-value < 0.05 and **** for p-value < 0.0001.

These results suggest that general skill learning mainly occur offline, but the performance increase during rest periods might be mainly due to fatigue or inhibition release (Rickard et al. 2008; Brawn et al. 2010; Török et al. 2017). To ensure that performance increase during rest periods reflects offline learning and not only fatigue/inhibition release, we analyzed rest periods following the first blocks of each session, during which no performance decrements were observed (average of the first blocks of the two sessions for Experiment 1, first block of the session for the Experiment 2 and average of the first blocks of the eight sessions for Experiment 3). Indeed, no decrease in performance occurred during these first session blocks; in Experiment 1 we even observed a modest performance increase (Experiment 1: *M_online_* = 11.13 ± 37.82 seconds, *t*(62) = 2.32, *p* = 0.02; Experiment 2: *M_online_* = 0.58 ± 30.25 seconds, *t*(179) = 0.26, *p* = 0.80; Experiment 3: *M_online_* = −0.53 ± 11.94 seconds, *t*(24) = 0.22, *p* = 0.83) but following rest periods were still accompanied by a general skill performance increase (Experiment 1: *M_offline_* = 43.44 ± 46.59 seconds, *t*(62) = 8.34, *p* < 10^−9^; Experiment 2: *M_offline_* = 27.77 ± 30.40 seconds, *t*(179) = 12.22, *p* < 10^−24^; Experiment 3: *M_offline_* = 20.29 ± 10.10 seconds, *t*(24) = 9.84, *p* < 10^−9^) (Fig. 2D, 2G, 2J).

In the ASRT tasks (Experiments 2 and 3), general skill learning over the task was also visible when considering only the *random-low* trials instead of all trials (Experiment 2: *r_s_* = −0.71, *p* < 10^−7^; Experiment 3: *r_s_* = −0.96, *p* < 10^−107^) and similar online and offline dynamics were found when considering only *random-low* trials when all blocks were included (Experiment 2: *M_online_* = −27.38 ± 10.93 seconds, *t*(179) = −33.51, *p* < 10^−78^ and *M_offline_* = 28.65 ± 11.08 seconds, *t*(179) = 34.58, *p* < 10^−80^; experiment 3: *M_online_* = −17.89 ± 6.38 seconds, *t*(24) = −13.74, *p* < 10^−12^ and *M_offline_* = 18.74 ± 6.24 seconds, *t*(24) = 14.70, *p* < 10^−12^) or when only the first blocks were included (Experiment 2: *M_online_* = 0.50 ± 52.60 seconds, *t*(177) = 0.12, *p* = 0.90 and *M_offline_* = 26.55 ± 57.41 seconds, *t*(177) = 6.15, *p* < 10^−8^; Experiment 3: *M_online_* = −3.92 ± 15.33 seconds, *t*(24) = 1.25, *p* = 0.22 and *M_offline_* = 18.74 ± 6.24 seconds, *t*(24) = 6.57, *p* < 10^−6^).

We also investigated whether offline general skill learning across day or week was also visible. In Experiment 1, offline change in general skill performance between sessions 12 hours apart was significant (*M_LongOffline_* = 28.23 ± 61.88 seconds, *t*(62) = 3.59, *p* < 10^−3^). In Experiment 3, offline change in general skill performance between sessions a week apart was not significant (*M_LongOffline_* = 5.00 ± 17.25 seconds, *t*(24) = 1.42, *p* = 0.17).

### Dynamics of statistical learning

Statistical learning, defined as the increase of the difference in reaction time between *random-high* and *random-low trials*, was present in both ASRT experiments (Experiment 2: *r_s_*= 0.81, *p* < 10^−10^; Experiment 3: *r_s_*= 0.72, p < 10^−32^). When looking at online *vs*. offline gain in performance, we observed that statistical learning increased during practice (Experiment 2: *M_online_* = 5.22 ± 15.27 seconds, *t*(179) = 4.58, *p* < 10^−5^; Experiment 3: *M_online_* = 7.37 ± 6.70 seconds, *t*(179) = 5.40, *p* < 10^−4^) and decreased during rest periods (Experiment 2: *M_offline_* = −5.06 ± 13.97 seconds, *t*(179) = −4.85, *p* < 10^−5^; Experiment 3: *M_offline_* = −5.05 ± 13.97 seconds, *t*(179) = 4.84, *p* < 10^−5^) (Fig. 3B, 3E). Statistical learning also decreased between sessions a week apart in Experiment 3 (*M_LongOffline_* = −20.00 ± 35.65 seconds, *t*(24) = −2.75, *p* < 0.02).

**Figure 3:**
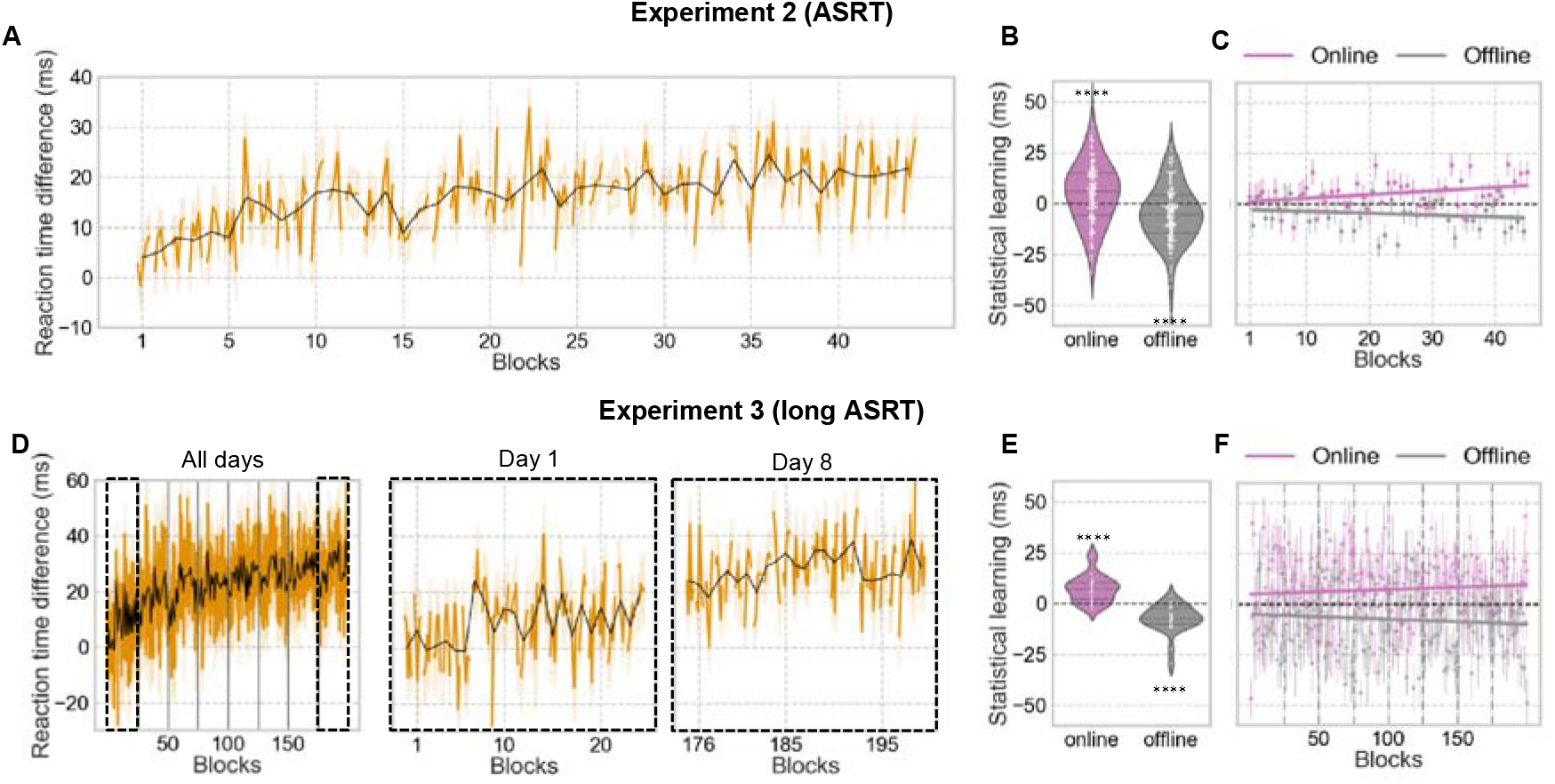
Statistical learning occurs during practice periods. (A) Average statistical learning (RT difference between random-high and random-low trials) per block (black line) and per bin (orange line) for Experiment 2 (ASRT). (B) Average online and offline statistical learning across all blocks for Experiment 2 (ASRT). (C) Online and offline statistical learning across all blocks and with a linear fit for Experiment 2. (D) Average statistical learning per block (black line) and per bin (orange line) for Experiment 3 (long ASRT). For better visualization, a zoom-in for day 1 and day 8 is represented. (E) Average online and offline statistical learning across all blocks with a linear fit for Experiment 3 (long ASRT). (F) Online and offline statistical learning across all blocks for Experiment 3 (long ASRT). Significance ****for p value<0.0001.

### Dynamics of high-order rule learning

High-order rule learning, defined as the increase of the difference in reaction time between *pattern* and *random-high trials*, was present only in Experiment 3 (long ASRT) (Experiment 2: *r_s_ = 0.17, p* = 0.27; Experiment 3: *r_s_* = 0.67, *p* < 10^−26^) (Fig. 4A, only Experiment 3 is displayed). When looking at online *vs*. offline gain in performance, we observed that high-order rule learning decreased during practice (Experiment 3: *M_online_* = −3.03 ± 6.15 seconds, *t*(24) = −2.42, *p* = 0.02) and increased during rest periods (Experiment 3: *M_offline_* = 2.76 ± 6.01 seconds, *t*(24) = 2.25, *p* = 0.03) (Fig. 4B). Change in high-order rule learning between sessions a week apart in Experiment 3 was not significant (*M_LongOffline_* = 6.72 ± 22.75 seconds, *t*(24) = 1.44, *p* = 0.16).

**Figure 4:**
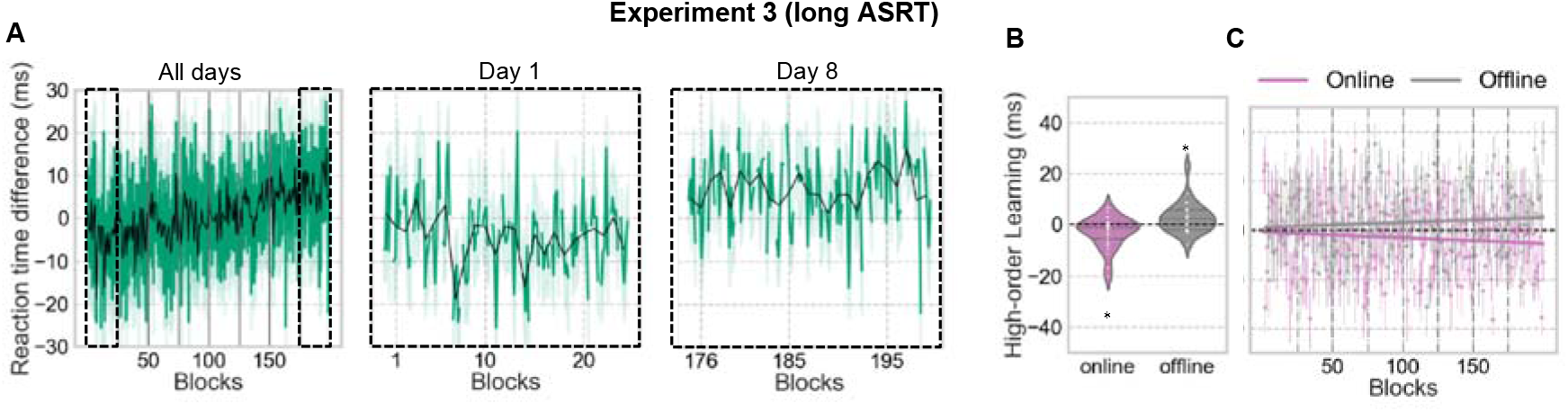
High-order rule learning occurs during rest periods. (A) Average high-order rule learning (RT difference between pattern and random-high trials) per block (black line) and per bin (green line) for Experiment 3 (long ASRT). For better visualization, a zoom-in for day 1 and day 8 is represented. (B) Average online and offline high-order rule learning across all blocks for Experiment 3 (long ASRT). (C) Online and offline high-order rule learning across all blocks and with a linear fit for Experiment 3. Significance is noted * for p value<0.5

## Discussion

Our brains can learn new skills very quickly. But the short-scale dynamic of this learning, and in particular, whether the new skill can be learned online or offline, has only recently started to be investigated (Du et al. 2016; Bönstrup et al. 2019, 2020). Here, we used three different experiments (one SRT and two ASRT tasks) and a total of 506 behavioral sessions to characterize the online and offline contribution for three types of learning, namely general skill learning, statistical learning and high-order rule learning. Our results revealed that the short-scale dynamics of different types of learning are mirroring each other, building-up either during practice or during the following rest periods. Specifically, statistical learning is acquired during practice periods, while general skill and high-order rule learning are acquired during break periods.

Statistical learning refers to the process of extracting probabilistic structure from the environment (Romberg and Saffran 2010; Sherman and Turk-Browne 2020). In our ASRT tasks, statistical learning is evidenced by shorter reaction times during triplets that appear frequently (*random-high trials*) compared to triplets that appear less frequently (*random-low trials*) (Howard and Howard 1997). Performance in statistical learning increases during practice and decreases during rest periods (Fig. 3). These results suggest that statistical learning benefits from evidence accumulation developing during practice and does not consolidate but decays during rest periods. This observation may explain why no evidence for offline consolidation of statistical learning was found during 12-hour sleep or awake periods (Song et al. 2007; Nemeth et al. 2010).

Conversely, higher-order rule learning, evidenced by faster performance during *pattern*relative to *random-high trials* specifically increases offline during rest periods (Fig. 4). This type of learning is much lower in magnitude than statistical learning and becomes significant only after many trials or sessions, as in the third experiment. Indeed, while the probabilistic learning in the ASRT task is based on acquiring the statistics on low order, simple transitions, the high-order rule learning is, as indicated by its name, based on acquiring the deterministic rule on high-order, complex transitions, i.e., every other trial. A potential explanation for these opposite results for these two learning types is that statistical knowledge on simple transitions can be acquired under attentional distraction coming from the task itself of mapping visual cues with response keys. In contrast, *higher-order* rule learning could need more attentional resources and consequently occurs only between practice periods. It has indeed been shown during sequence learning that simple transitions (Jiménez and Vázquez 2005; Rowland and Shanks 2006; Nemeth et al. 2011), but not more complex structures (Cohen et al. 1990), could be learned under attentional distraction.

Another possible explanation stands in the deterministic *vs*. probabilistic nature of these two types of learning. While deterministic and probabilistic information may be considered as a continuum of the same process (deterministic rule is mathematically an extreme case of statistical information with probabilities of 0 or 1), past research suggests that both processes are linked to different brain regions (Bhanji et al. 2010), influenced differently by the explicitness of the information (Stefaniak et al. 2008) and better modeled by two distinct hypothesis spaces instead of one (Maheu et al. 2020). It is then possible that uncertain regularities (statistical learning) need evidence accumulation and can only be acquired online while deterministic regularities (rule learning) need a rest period to be consolidated, maybe because they are somehow rehearsed or replayed during rest. Future studies will have to dissociate whether this difference in dynamics between statistical and high-order rule learning is related to the low-order/high-order or the probabilistic/deterministic nature of the learning, or a mixture of both.

Our results also demonstrate that general skill learning is acquired during rest periods (Fig 2). This result stands both when the measure for general skill learning included all trials or only *random-low* trials, excluding then any predictable patterns from the stimulus stream. It thus suggests that the fast-consolidation of procedural learning during breaks observed in previous research (Du et al. 2016; Bönstrup et al. 2019, 2020) may be less dependent of the sequence learning itself but depends more on a mixture of improvement in sensorimotor transformation, dexterity, and familiarization with the task. While statistical and high-order rule learning are measured as a difference between two types of trials, precluding that the offline gap in performance is due to a release of fatigue or reactive inhibition effect (Török et al. 2017), the general skill learning is measure by a reaction time, which is very sensitive to fatigue, as depicted by the constant decrease in reaction time within blocks in the three experiments (Fig. 2A, E, H). To ensure that the offline gap in general skill performance is not simply a release of fatigue, we tested the offline change in general skill performance after the first blocks of each session during which there is no decrease in reaction time (Fig. D, G, J) and the offline gain was still present. Offline improvements in general skills are thus not only related to fatigue release but are likely to also reflect consolidation processes (see also Bönstrup et al. 2020).

In this study, we identified the short-scale temporal dynamics of three types of learning, namely general skill learning, statistical learning and high-order rule learning, extracted from the same information stream. We revealed that they are not developing at the same time, with general skill and high-order rule learning developing offline while statistical learning is developing online. These results suggest that such types of learning rely on separate neural mechanisms with their own dynamics. Our unprecedented dissection of the short-scale dynamics of subcomponents of learning challenge the classical view of memory acquisition and consolidation, which would be applied indifferently to all types of learning. We revealed, on the contrary, that statistical learning occurs only during practice and general skill and high-order rule learning occur only during breaks.

## Methods

### Participants

Two hundred and sixty-eight (268) healthy young volunteers participated in three studies (192 women, 76 men, mean age = 22.2 years) for a total of 506 reported behavioral sessions. All participants had normal or corrected-to-normal vision, and none of them reported a history of any neurological and/or psychiatric condition. Participants provided informed consent to the procedure before enrollment, as approved by the institutional review board of the local research ethics committee. The three experiments were approved by the United Ethical Review Committee for Research in Psychology (EPKEB) in Hungary and by the research ethics committee of Eötvös Loránd University, Budapest, Hungary. The experiments were conducted in accordance with the Declaration of Helsinki. Participants received course credits for taking part in the experiment. Data from Experiment 2 were previously published (Kóbor et al. 2017; Török et al. 2017). The results of the present paper were not tested nor reported before. Figure. 1 summarizes the design of the present study.

### Serial Reaction Time (SRT) Task

During the Serial Reaction Time (SRT) task (Nissen and Bullemer 1987), four empty circles were horizontally arranged on the screen. Participants were instructed to respond to a stimulus (a dog’s head) that appeared in one of the four open circles by pressing one of four corresponding keys on a computer keyboard (Z, C, B, or M on a QWERTY keyboard) as quickly and accurately as possible after the appearance of the stimulus. Participants used their left and right middle and index fingers to respond to the stimuli. The stimulus remained visible until participants pressed the correct key, at which time it disappeared. The following stimulus appeared 120 ms after the offset of the previous stimulus. The SRT task was programmed and displayed using E-prime software (Psychology Software Tools, Inc.). The serial order of the four possible positions (coded as 1, 2, 3, and 4) in which target stimuli could appear was determined by a twelve-element sequence (2-3-1-4-3-2-4-1-3-4-2-1) (Robertson 2007). An experimental session was divided into blocks with either 60 trials corresponding to five repetition of the twelve-element sequence or 60 pseudo-random trials in which the visual cue no longer played out a deterministic pattern of positions.

### Alternating Serial Reaction Time (ASRT) Task

The visual display, response modality, timing, instructions, and program software for the ASRT task were similar to those during the SRT task. The serial order of the four possible positions (coded as 1, 2, 3, and 4) in which target stimuli could appear was determined by an eight-element sequence (Howard and Howard 1997; Song et al. 2007; Janacsek et al. 2012). In this sequence, every second element appeared in the same order during the entire task, while the other elements’ positions were randomly chosen (e.g., 2-*r*-1-*r*-3-*r*-4-*r*, where numbers refer to a predetermined location in one of the four locations and *r* refer to randomly chosen locations out of the four possible). A total of six unique sequences of predetermined elements were created and one of them was assigned to each subject in a random order (Howard and Howard 1997). An experimental session was divided into blocks starting with five random trials (warm-up) followed by the eight-element sequence repeated ten times (Nemeth et al. 2010; Nemeth, Janacsek, Király, et al. 2013). Warm-up trials were discarded from the analyses.

Due to the alternating sequence structure, some patterns of three consecutive elements (henceforth referred to as triplets) occurred with a higher probability than other ones. Each trial was categorized as the last element of either a high- or a low-probability triplet. High-probability triplets could be formed either by predetermined elements or random ones. In the above sequence example (2-*r*-1-*r*-3-*r*-4-*r*), the probability that a triplet starting with the element ‘2’ and ending with the element ‘1’ occurred was of 62.5%. Indeed, the item ‘2’ could be either predetermined (50%) or random (50%). If it is predetermined, then the last element of the triplet has to be ‘1’; if it is random the last element of the triplet could be anything. Thus, the item ‘1’ had 50% probability of occurring as the last predetermined element of the triplet plus 12.5% of chances to occur as a random element. In contrast, triplets such as 1-x-2 or 4-x-3 occurred with a low probability (12.5%) because they could only occur when the third element of the triplet was random. Low-probability triplets forming repetitions (e.g., 222) or trills (e.g., 232) were discarded from analyses as participants often show preexisting response tendencies to them (Howard et al. 2004; Soetens et al. 2004). Trials were participants pressed a wrong button were also discarded. Participants were not informed of any regularity. Each trial could be a *pattern trial, a random-high trial or a random-low trial*. A *pattern trial* corresponded to a predetermined element ending a triplet (all pattern trials are high-probability triplets); a *random-high trial* corresponded to a random element ending a high-probability triplet; a *random-low trial* corresponded to a random element ending a low-probability triplet. This sequence structure allows the distinction between (i) general skill learning, measured by a decrease in reaction time (RT) for all trials, (ii) statistical learning, measured by the difference in RT between the *random-high trials* and the *random-low trials* (because they end two types of triplets that appear randomly, but *random-high trials* are more frequent than *random-low trials*) and (iii) high-order deterministic learning, measured by the difference in RT between *pattern trials* and *random-high trials* (because they end two types of triplets that are similar in term of sequence but *pattern trials*, unlike *random-high trials*, are predictable) (Howard and Howard 1997; Nemeth, Janacsek, and Fiser 2013).

### Procedure: Experiment 1

Sixty-three participants took part in this experiment. They each performed two sessions separated by 12 hours. Each session contained a total of 13 blocks of SRT task, with the 6^th^ and the 12^th^ block displaying random sequences. Behavioral performances during random blocks were discarded from the analyses (but there are visible in Fig. 2A for illustration purpose). After each block, the average speed and accuracy for the most recent block were displayed to the participants, and they could have a short break before starting the next block by pressing a button. The average block duration across participants and blocks was 31.33 ± 5.11 seconds. The average break duration across participants and breaks was 24.26 ± 19.83 seconds.

### Procedure: Experiment 2

One hundred eighty participants took part in this experiment. They each performed one session of 45 blocks of ASRT task. After each block, the average speed and accuracy for the most recent block were displayed to the participants, and they could have a short break before starting the next block by pressing a button. After 15 blocks and 30 blocks, participants had a more extended break and filled questionnaires. The average block duration across participants and blocks was 46.45 ± 3.34 seconds. The average short break duration across participants and blocks was 18.75 ± 10.7 seconds. The average break duration for the two longer breaks with questionnaire was 258.0 ± 99.75 seconds.

### Procedure: Experiment 3

Twenty-five participants took part in this experiment. They each performed eight sessions of 25 blocks of ASRT task. Each session was a week apart. After each block, the average speed and accuracy for the most recent block were displayed to the participants, and they could have a short break before starting the next block by pressing a button. The average block duration across participants and blocks was 41.79 ± 3.78 seconds. The average break duration across participants and breaks was 18.56 ± 3.31 seconds.

### Learning measures and statistical analyses

General skill learning was defined as a decrease of RT for all trials across blocks. In ASRT tasks, general skill learning was also tested considering random low trials only. Statistical and high-order rule learning was measurable only in ASRT experiments. Statistical learning was defined as an increase of RT difference between *random-low* and *random-high trials* (RT_random-low_-RT_random-high_) across blocks. Higher-order rule learning was defined as an increase of RT difference between *random-high* and *pattern trials* (RTrandom-high-RTpattern). High-order rule learning takes a high number of trials or sessions in ASRT to become visible. Indeed, in the current study, it was only observable in the long ASRT task (Experiment 3, see Results section). To estimate general skill, statistical and high-order rule learning over the tasks, Spearman correlation between learning measures (block-average RT for general skill learning or block-average difference in RT between two types of triplet for statistical and higher-order deterministic learning) and block position was used. To measure the online (over practice blocks) and offline (over rest periods) contribution to each type of learning, in both SRT and ASRT tasks, each block was binned into five bins. Each bin corresponds to 12 trials (one 12-element sequence) in the SRT task and 16 trials (two 8-element sequences) in the ASRT task. Online learning was measured as the difference in learning between the last bin of a block and the first bin of the same block. Offline learning was measured as the difference in learning between the first bin of a block and the last bin of the previous block (Fig. 2B). For general skill learning, as learning is defined as a decrease in RT, online and offline measures were reversed so that learning appears positive on the violin plots (Fig. 2C). One-sample t-tests against zero were used to assess if learning occurred during practice (online) or rest (offline) periods.

